# Predicting protein stability changes upon mutations with dual-view ensemble learning from single sequence

**DOI:** 10.1101/2024.04.22.590665

**Authors:** Zhiwei Nie, Yiming Ma, Yutian Liu, Xiansong Huang, Zhihong Liu, Peng Yang, Fan Xu, Feng Yin, Zigang Li, Jie Fu, Zhixiang Ren, Wen-Bin Zhang, Jie Chen

## Abstract

Predicting the protein stability changes upon mutations is one of the effective ways to improve the efficiency of protein engineering. Here, we propose a dual-view ensemble learning-based framework, DVE-stability, for mutation-induced protein stability change prediction from single sequence. DVE-stability integrates the global and local dependencies of mutations to capture the intramolecular interactions from two views through ensemble learning, in which a structural microenvironment simulation module is designed to indirectly introduce the information of structural microenvironment at the sequence level. DVE-stability achieved state-of-the-art prediction performance on 7 single-point mutation benchmark datasets, and comprehensively surpassed other methods on 5 of them. Furthermore, DVE-stability outperformed other methods comprehensively through zero-shot inference on multiple-point mutation prediction task, demonstrating superior model generalizability to capture the epistasis of multiple-point mutations. More importantly, DVE-stability exhibited superior generalization performance in predicting rare beneficial mutations that are crucial for practical protein directed evolution scenarios. In addition, DVE-stability identified important intramolecular interactions via attention scores, demonstrating interpretable. Overall, DVE-stability provides a flexible and efficient tool for mutation-induced protein stability change prediction in an interpretable ensemble learning manner.

## 1 Introduction

Quantifying the mutation-induced effects is fundamental for exploring the evolutionary fitness landscape of proteins [1, 2]. Protein thermodynamic stability is of particular interest as it provides valuable insights into protein folding and functions [3, 4], thereby promoting the protein engineering [5–8] for pharmaceutical biologics and industrial biocatalysts. Considering that traditional experimental measurements of protein stability changes are costly in both time and money, computational methods for predicting protein stability changes upon mutations have emerged in an endless stream [9–12].

Existing computational methods for mutation-induced protein stability change prediction, especially deep-learning approaches, can be divided into two categories: structure-based and sequence-based methods. The structure-based methods predict stability changes for protein point mutations through protein structure input [13– 21]. ThermoMPNN [19] employed a pretrained ProteinMPNN model [22] for protein structure encoding, which was used for subsequent stability change prediction module. Stability Oracle [20] pretrained a graph-transformer backbone for stability change regression prediction. GeoDDG-3D [21] utilized a pretrained geometric encoder of graph attention neural network architecture. In contrast, sequence-based methods predict stability changes upon mutations given only protein sequence [23, 23–29]. Mutate Everything [28] utilized a pretrained feature extractor (ESM[30] or AlphaFold[31]) to compute amino-acid-level features of the input sequence. PROSTATA [27] employed pretrained ESM models as the embedding backbone of protein sequences. SPIRED-Stab [29] developed a pretrained single-sequence-based structure prediction model for protein stability change prediction. For structure-based methods, a small number of mutations are unlikely to cause significant changes in the three-dimensional structure, so that the performance of structure-based methods is limited to extremely similar structural features. Sequence-based methods can obtain sequence features with greater discrimination, but they cannot encode the structural microenvironment in which the mutation is located, which makes their generalization ability encounter bottlenecks.

In the protein engineering scenario of a real laboratory, the performance of a predictive method determines the engineering speed, while the generalization ability of a predictive method determines its scope of application. Therefore, to take into account both prediction performance and generalization ability, we propose a **D**ual-**V**iew **E**nsemble-learning-based framework, DVE-stability, for protein stability change prediction based on protein sequence. DVE-stability integrates the global dependencies of mutations dominated by long-range interactions and local dependencies of mutations determined by the structural microenvironments, enabling the neural network to capture the intramolecular interaction network in which the mutation is located from two views through ensemble learning. Considering the importance of the structural microenvironments for stability changes caused by mutations, we argue that it is beneficial to introduce information about the structural microenvironment into sequence-based methods in an indirect way. To this end, we propose the structural microenvironment simulation (SMS) module to embed in the DVE-stability framework to indirectly introduce the information of structural microenvironment at the sequence level. DVE-stability achieved state-of-the-art prediction performance on 7 single-point mutation benchmark datasets, and comprehensively surpassed other methods on 5 of them. More importantly, DVE-stability exhibited superior generalization performance through zero-shot inference on multiple-point mutation prediction task and rare beneficial mutation mining task. Overall, DVE-stability represents an interpretable ensemble learning approach for protein stability change prediction upon mutations from single sequence, which is expected to provide an efficient and generalizable tool for protein engineering.

## 2 Results

### 2.1 The dual-view ensemble learning architecture of DVE-stability

The dual-view ensemble learning architecture of DVE-stability (Fig.1a) is designed to simultaneously internalize the global dependencies of mutations dominated by longrange interactions and local dependencies of mutations determined by the structural microenvironments. First, a mutated protein sequence and its wild-type sequence are encoded in parallel by the pretrained protein language model (PLM), ESM-2-650M [30], to extract corresponding amino-acid-level embeddings. Second, the dual-view interaction dependencies of mutations are extracted in the mutated protein, as well as in the wild-type protein. It is worth noting that during the training process, the employed ESM-2-650M is directly fine-tuned with all parameters. The CLS token is a special marker added to the beginning of the input sequence in Transformer-based architectures [32], and its output vector is designed to be the global semantic representation of the entire input sequence. In tasks such as text classification [33], semantic matching [34], and contrastive learning [35], it carries the abstraction of the meaning of the entire sentence, making it easy to use directly in downstream tasks.

**Fig. 1.**
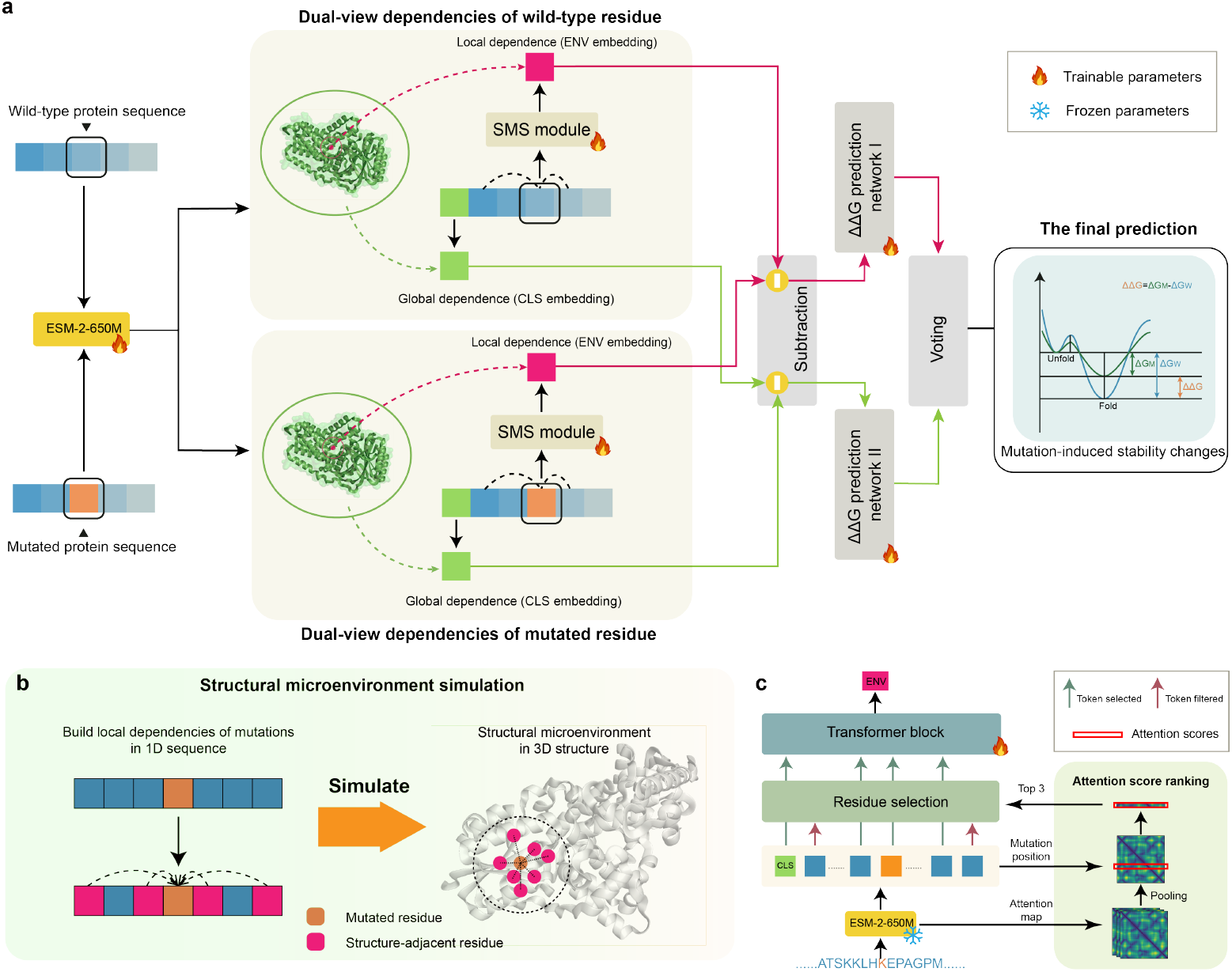
The model architecture of DVE-stability. **a**, The architecture overview of DVE-stability. The wild-type and mutated protein sequences are input in parallel to extract dual-view interaction dependencies of mutations (CLS embedding and ENV embedding) for subsequent ensemble learning. **b**, Illustration of structural microenvironment simulation (SMS) module, where residues structurally adjacent to the mutated residue are identified in the protein sequence to build the local dependencies of the mutation, thereby simulating the structural microenvironment in 3D structure. **c**, Internal architecture of the SMS module. The off-the-shelf protein language model ESM-2-650M is employed to obtain the ranking of the attention scores of the residue types at the mutation position, where the residues with the top 3 largest attention scores are selected. A transformer block is adopted to fuse the above selected residues to obtain an “ENV” embedding that represents local dependencies of mutations. **Alt text**: A three-panel schematic illustrating the DVE-stability framework, the structural microenvironment simulation module, and the internal architecture of this module.

In this work, the CLS token is employed to represent the global dependencies of mutations on the entire protein sequence, while the local dependencies of mutations on its structurally adjacent residues is built through a structural microenvironment simulation (SMS) module. As shown in Fig.1b, considering that fitness effects upon mutations are largely determined by the structural microenvironment of mutations [36–38], we pursue the simulation of structural microenvironment at the sequence level, thereby indirectly constructing and encoding the local dependencies of mutations in 3D structures. To this end, the SMS module is designed to identify the related residues that may constitute the structural microenvironment of mutations and perform feature fusion. Specifically, considering that the attention map of PLM has been proven to capture residue contact relationships to a certain extent [39, 40], ESM-2-650M with frozen parameters is adopted for the identification of structurally adjacent residues based on the assigned attention scores. The top 3 residues with the highest attention scores are selected, while the remaining residues are filtered out. Subsequently, the embeddings of selected residues are fused through a transformer block into an “ENV” embedding that represents local dependencies of mutations (Fig.1c).

Third, the global dependencies of the mutation in a mutated protein sequence is subtracted by that in its wild-type sequence to highlight the mutational effects on the intramolecular interaction network of proteins, and the local dependencies of the mutation are also processed in the same way. Fourth, the difference between global dependencies and local dependencies is used for the prediction of protein stability changes upon mutations (i.e. ΔΔ*G*) separately, and the final prediction is the average of the two branch predictions.

### 2.2 Prediction performance for single-point mutations

To evaluate the prediction performance of DVE-stability for protein stability changes upon single-point mutations, the widely recognized benchmark dataset S669 [9] is adopted for the test set. Following the evaluation setting (trained with high-throughput dataset) of a prior work Mutate Everything [28], we compared the prediction performance of different methods for single-point mutations in Table 1. It is worth noting that we created a reverse dataset of S669 to ensure the robustness of the evaluation (Methods 4.1.1). Through comprehensive comparisons of classification (stabilizing mutation or not) and regression (ΔΔ*G* regression) prediction tasks, we found that all evaluation metrics of DVE-stability were the best or sub-optimal compared with other prediction methods, reaching the state-of-the-art prediction performance. Furthermore, to ensure the robustness of the evaluation, we evaluated the prediction performance of different methods on another 6 single-point mutation datasets, namely S461, S783, Myoglobin, PTEN&TPMT, S349, and T2837. The prediction results can be seen in Supplementary Table S1-S4, where our DVE-stability model achieved state-of-the-art prediction performance on all the additional datasets.

**Table 1.** Prediction performance comparison for single-point mutations on direct and reverse S669 datasets. Bold values represent the best performance, and underlined values represent suboptimal performance. The raw results of the other models are taken from a prior work Mutate Everything [28].

| Method | S669 direct |  |  |  | S669 reverse |  |  |  |
| --- | --- | --- | --- | --- | --- | --- | --- | --- |
| | $\rho$ | AUC | MCC | RMSE | $\rho$ | AUC | MCC | RMSE |
| Structure-based |  |  |  |  |  |  |  |  |
| mCSM | 0.37 | 0.66 | 0.13 | 1.53 | 0.24 | 0.61 | 0.10 | 2.33 |
| I-Mutant3.0 | 0.35 | 0.64 | 0.08 | 1.53 | 0.17 | 0.59 | 0.06 | 2.35 |
| DUET | 0.42 | 0.68 | 0.19 | 1.52 | 0.26 | 0.63 | 0.12 | 2.16 |
| FoldX | 0.27 | 0.62 | 0.14 | 2.35 | 0.32 | 0.67 | 0.20 | 2.54 |
| MAESTRO | 0.46 | 0.69 | 0.26 | 1.45 | 0.22 | 0.61 | 0.11 | 2.12 |
| PopMusic | 0.42 | 0.69 | 0.22 | 1.51 | 0.24 | 0.62 | 0.12 | 2.10 |
| SDM | 0.39 | 0.67 | 0.21 | 1.67 | 0.14 | 0.59 | 0.12 | 2.15 |
| INPS3D | 0.44 | 0.70 | 0.20 | 1.49 | 0.36 | 0.69 | 0.23 | 1.78 |
| Dynamut | 0.38 | 0.68 | 0.20 | 1.59 | 0.37 | 0.67 | 0.17 | 1.69 |
| ThermoNet | 0.38 | 0.69 | 0.21 | 1.62 | 0.35 | 0.66 | 0.18 | 1.66 |
| PremPS | 0.42 | 0.66 | 0.20 | 1.50 | 0.43 | 0.66 | 0.22 | 1.49 |
| DDGun3D | 0.43 | 0.71 | 0.27 | 1.60 | 0.41 | 0.71 | 0.23 | 1.61 |
| ACDC-NN | 0.46 | 0.73 | 0.23 | 1.49 | 0.45 | 0.72 | 0.22 | 1.50 |
| Stability Oracle | 0.53 | 0.75 | 0.34 | 1.44 | 0.53 | <u>0.75</u> | **0.32** | **1.43** |
| Sequence-based |  |  |  |  |  |  |  |  |
| MuPro | 0.27 | 0.61 | 0.03 | 1.60 | 0.22 | 0.64 | 0.08 | 2.41 |
| I-Mutant3.0-Seq | 0.34 | 0.67 | 0.25 | 1.54 | 0.23 | 0.61 | 0.05 | 2.25 |
| DDGun | 0.43 | 0.72 | 0.24 | 1.73 | 0.41 | 0.71 | 0.23 | 1.76 |
| ACDC-NN-Seq | 0.44 | 0.72 | 0.26 | 1.52 | 0.44 | 0.72 | 0.23 | 1.52 |
| INPS-Seq | 0.44 | 0.72 | 0.25 | 1.52 | 0.44 | 0.72 | 0.21 | 1.53 |
| PROSTATA | 0.50 | 0.73 | 0.28 | 1.44 | <u>0.50</u> | 0.73 | 0.29 | 1.44 |
| Mutate Everything | <u>0.56</u> | <u>0.76</u> | **0.35** | <u>1.38</u> | 0.49 | 0.73 | 0.30 | 1.48 |
| DVE-stability | **0.59** | **0.78** | <u>0.34</u> | **1.35** | **0.54** | **0.76** | <u>0.30</u> | <u>1.44</u> |

In conventional evaluation protocol, the evaluation metrics are calculated using a cross-protein evaluation method (i.e., for all protein mutants in the test set). This evaluation strategy is insufficient to reflect the real ability of the predictive model to rank multiple single-point mutations of a single protein, which is more relevant in practical protein engineering scenario. Therefore, we followed the evaluation protocol of a prior work SPIRED-Stab [29], where the internal metrics of each protein were first evaluated, and then the average and standard deviation of each metric in all tested proteins were reported. As shown in Table 2, all evaluation metrics (namely Spearman, Kendall and Pearson correlation coefficients, and Top-K precision) of DVE-stability trained with S2648 were the best or sub-optimal compared with other prediction methods, demonstrating its applicability for practical protein engineering.

**Table 2.** Comparison of single-point mutation ranking performance on S669 dataset. Bold values represent the best performance, and underlined values represent suboptimal performance. The raw results of the other models are taken from a prior work SPIℝED-Stab [29]. *ρ, τ*, and *r* denote Spearman, Kendall and Pearson correlation coefficients, respectively. Top-5 and Top-10 denote the Top-5 precision and Top-10 precision, respectively.

| Method | $\rho$ | $\tau$ | $r$ | Top-5 | Top-10 |
| --- | --- | --- | --- | --- | --- |
| **Structure-based** |  |  |  |  |  |
| SDM | 0.31± 0.33 | 0.22 ± 0.26 | 0.33 ± 0.36 | 0.33 ± 0.22 | 0.57 ± 0.23 |
| ThermoNet | 0.31 ± 0.30 | 0.22 ± 0.23 | 0.38 ± 0.30 | 0.31 ± 0.18 | 0.54 ± 0.24 |
| mCSM | 0.33 ± 0.30 | 0.23 ± 0.23 | 0.35 ± 0.31 | 0.39 ± 0.14 | 0.53 ± 0.23 |
| Dynamut | 0.34 ± 0.31 | 0.25 ± 0.23 | 0.34 ± 0.32 | 0.32 ± 0.27 | 0.52 ± 0.27 |
| I-Mutant3.0 | 0.34 ± 0.30 | 0.26 ± 0.22 | 0.39 ± 0.29 | 0.39 ± 0.18 | 0.54± 0.23 |
| MAESTRO | 0.35 ± 0.33 | 0.26 ± 0.25 | 0.40 ± 0.33 | 0.37 ± 0.13 | 0.59 ± 0.19 |
| DUET | 0.38 ± 0.30 | 0.27 ± 0.24 | 0.41 ± 0.30 | 0.40 ± 0.17 | 0.56 ± 0.24 |
| ACDC-NN | 0.40 ± 0.30 | 0.30 ± 0.23 | 0.42 ± 0.33 | 0.44 ± 0.20 | 0.57 ± 0.26 |
| FoldX | 0.41 ± 0.35 | 0.31 ± 0.28 | 0.36 ± 0.39 | 0.36 ± 0.29 | 0.59 ± 0.25 |
| PoPMuSiC | 0.42 ± 0.29 | 0.30 ± 0.23 | 0.45 ± 0.30 | 0.37 ± 0.23 | 0.63 ± 0.20 |
| DDGun3D | 0.43 ± 0.31 | 0.32 ± 0.24 | 0.48 ± 0.29 | 0.41 ± 0.21 | 0.62 ± 0.21 |
| PremPS | 0.45 ± 0.29 | 0.33 ± 0.24 | 0.48 ± 0.28 | 0.39 ± 0.21 | 0.58 ± 0.27 |
| DDMut | 0.46 ± 0.37 | 0.35 ± 0.29 | 0.52 ± 0.28 | 0.44 ± 0.26 | 0.59 ± 0.20 |
| INPS3D | 0.46 ± 0.27 | 0.35 ± 0.21 | 0.48 ± 0.28 | 0.44 ± 0.22 | 0.59 ± 0.22 |
| GeoDDG-3D | 0.50 ± 0.28 | 0.38 ± 0.22 | 0.51 ± 0.27 | 0.53 ± 0.22 | 0.69 ± 0.16 |
| RaSP | 0.52 ± 0.29 | 0.39± 0.23 | 0.50 ± 0.28 | 0.45 ± 0.23 | 0.62 ± 0.24 |
| Pythia | 0.55 ± 0.30 | 0.41 ± 0.22 | 0.55 ± 0.30 | 0.49 ± 0.26 | 0.64 ± 0.18 |
| ThermoMPNN | 0.56 ± 0.31 | 0.42 ± 0.25 | 0.53 ± 0.31 | 0.48 ± 0.20 | 0.63 ± 0.20 |
| **Sequence-based** |  |  |  |  |  |
| MUPro | 0.18 ± 0.33 | 0.13 ± 0.23 | 0.21 ± 0.34 | 0.31 ± 0.24 | 0.53 ± 0.25 |
| I-Mutant3.0-Seq | 0.22 ± 0.37 | 0.16 ± 0.27 | 0.28 ± 0.33 | 0.33 ± 0.22 | 0.51 ± 0.23 |
| SAAFEC-SEQ | 0.34 ± 0.30 | 0.26 ± 0.22 | 0.39 ± 0.29 | 0.39± 0.18 | 0.54 ± 0.23 |
| ACDC-NN-Seq | 0.39 ± 0.29 | 0.29 ± 0.21 | 0.44 ± 0.29 | 0.43 ± 0.15 | 0.58 ± 0.21 |
| INPS-Seq | 0.40 ± 0.28 | 0.28± 0.22 | 0.44 ± 0.26 | 0.47 ± 0.20 | 0.56 ± 0.25 |
| DDGun | 0.45 ± 0.27 | 0.34 ± 0.21 | 0.47 ± 0.27 | 0.44 ± 0.20 | 0.56 ± 0.25 |
| PROSTATA | 0.50 ± 0.28 | 0.36±0.22 | 0.53± 0.27 | 0.49 ± 0.26 | 0.62±0.26 |
| Mutate everything (ESM-2) | 0.50 ± 0.34 | 0.38 ± 0.25 | 0.52 ± 0.30 | 0.41 ± 0.26 | 0.62 ± 0.23 |
| GeoDDG-Seq | 0.51 ± 0.28 | 0.39 ± 0.22 | 0.52 ± 0.29 | 0.49± 0.25 | 0.63 ± 0.22 |
| SPIRED-Fitness (zero-shot) | 0.54 ± 0.25 | 0.41± 0.20 | 0.52 ± 0.27 | 0.51 ± 0.18 | 0.64± 0.22 |
| GeoStab v2 | 0.55 ± 0.41 | 0.43 ± 0.32 | 0.54± 0.41 | 0.52 ± 0.26 | **0.69** ± 0.22 |
| SPIRED-Stab | 0.58 ± 0.34 | **0.46** ± 0.27 | 0.55 ± 0.36 | **0.59**± 0.22 | 0.67± 0.22 |
| DVE-stability | **0.59**± 0.29 | <u>0.43</u>± 0.33 | **0.55**± 0.31 | 0.54 ± 0.28 | 0.68 ± 25 |

### 2.3 Generalization performance for multiple-point mutations

DVE-stability can be generalized to predict stability changes upon multiple-point mutations through zero-shot inference. When input multiple-mutated proteins, it is only necessary to average the ENV features of multiple mutations and then perform ensemble learning with CLS. Following the evaluation settings of a prior work GeoDDG [21], we only use single-point mutation data (S8754 dataset) to train DVE-stability and then test it on the M1261 dataset (792 pieces of double mutation data and 469 pieces of triple or higher-order mutation data). As shown in Table 3, DVE-stability surpassed other methods comprehensively through zero-shot inference, whether on double mutation data or triple or higher-order mutation data. It is worth noting that even though DynaMut2 [41] and DDMut [42] were trained using multiple-point mutation data and GeoDDG-3D reutilized a model pretrained on a large amount of fitness data from deep mutational scanning, DVE-stability still outperformed them comprehensively, demonstrating a superior model generalizability to capture the epistasis of multiple-point mutations.

**Table 3.**
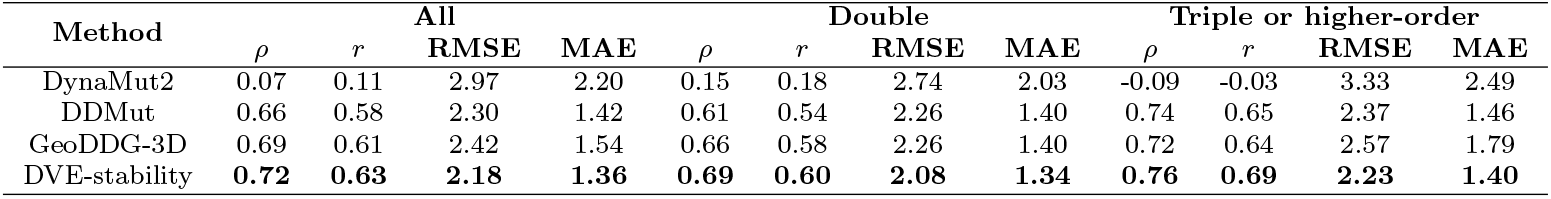
Prediction performance comparison for multiple-point mutations on M1261 dataset. Bold values represent the best performance. The raw results of the other models are taken from a prior work GeoDDG [21].

### 2.4 Generalization performance for rare beneficial mutations

In the vast evolutionary space of proteins, directed evolution in the laboratory needs to seek rare beneficial mutations that can improve protein fitness [43]. Considering that mutations that are beneficial to fitness improvement constitute a rare subset of the fitness landscape for protein evolution [44], accurately mining rare beneficial mutations is one of the potential means to improve the efficiency of directed evolution of proteins. This zero-shot prediction task seeks to avoid missing rare beneficial mutations, that is, as high a “recall” metric as possible.

To compare the generalization performance of different prediction methods on the real wet-lab directed evolution scenario, we selected a representative enzyme engineering case (*Is*PETase engineering with improved thermal stability [45]) to set up a validation experiment. In this validation experiment, there were 85 single mutations, including 21 beneficial mutations and 64 harmful mutations (Supplementary Table S5-S6). Three types of recent high-performance methods were adopted for prediction performance comparison, namely unsupervised protein evolution methods (PLM ensemble voting strategy, referred to as PLMVoting), structure-based supervised ΔΔ*G* prediction methods (ThermoMPNN [19], GeoDDG-3D [21]), and sequence-based supervised ΔΔ*G* prediction methods (PROSTATA [27], Mutate Everything [28], SPIRED-Stab [29]).

To ensure statistical robustness, the bootstrapping approach was adopted to construct 5,000 pseudo datasets, and then the mean, standard deviation and confidence interval of the recall metric of the above 5,000 experiments were reported. The prediction results of different methods are shown in Table 4, where we can find that DVE-stability surpassed other predictive methods. Overall, DVE-stability achieved superior generalization performance in mining rare beneficial mutations that are critical for practical protein engineering.

**Table 4.** Generalization performance of different methods for rare beneficial mutation mining. A bootstrapping approach was adopted to construct 5,000 pseudo datasets, and the most important metric “recall” for practical protein directed evolution was reported. Bold values represent the best performance.

| Method | Recall |  |
| --- | --- | --- |
|  | Mean/STD | 90% Confidence Interval |
| **Unsupervised** |  |  |
| PLMVoting | 0.38±0.147 | [0.20, 0.57] |
| **Structure-based supervised** |  |  |
| ThermoMPNN | 0.28±0.132 | [0.10, 0.45] |
| GeoDDG-3D | 0.53±0.147 | [0.11, 0.45] |
| **Sequence-based supervised** |  |  |
| PROSTATA | 0.57±0.146 | [0.38, 0.75] |
| Mutate Everything | 0.43±0.146 | [0.25, 0.61] |
| SPIRED-Stab | 0.53±0.146 | [0.34, 0.72] |
| DVE-stability | **0.67±0.139** | [0.50, 0.85] |

### 2.5 Ablation study

To evaluate the impact of key components on the prediction performance of DVE-stability, we conducted multiple ablation experiments, as shown in Table 5. To quantify the impact of three key components (ensemble learning, CLS embedding, ENV embedding), we built four model variants of DVE-stability, namely DVE-stability without ensemble learning (w/o ensemble), DVE-stability without CLS embedding (w/o CLS), DVE-stability without ENV embedding (w/o ENV), and DVE-stability without all components (w/o all). Whether on the direct or reverse dataset of S669, the removal of any component led to a decrease in prediction performance, among which the removal of ENV embedding led to the most significant performance degradation. When all key components were removed, the prediction performance of DVE-stability further deteriorated. In summary, the three key components are beneficial to the predictive ability of DVE-stability.

**Table 5.** Ablation experiments for key components. Bold values represent the best performance.

| Method | S669 direct |  |  |  | S669 reverse |  |  |  |
| --- | --- | --- | --- | --- | --- | --- | --- | --- |
| | $r\uparrow$ | AUC↑ | MCC↑ | RMSE↓ | $r\uparrow$ | AUC↑ | MCC↑ | RMSE↓ |
| DVE-stability | **0.59** | **0.78** | **0.34** | **1.35** | **0.54** | **0.76** | **0.30** | **1.44** |
| w/o ensemble | 0.52 | 0.75 | 0.30 | 1.46 | 0.51 | 0.74 | 0.29 | 1.47 |
| w/o ENV | 0.47 | 0.70 | 0.27 | 1.50 | 0.45 | 0.70 | 0.26 | 1.52 |
| w/o CLS | 0.58 | 0.77 | 0.32 | 1.40 | 0.52 | 0.76 | 0.30 | 1.45 |
| w/o all | 0.47 | 0.70 | 0.26 | 1.51 | 0.45 | 0.70 | 0.26 | 1.53 |

In addition, we also explored the impact of different types of training data on DVE-stability. Specifically, we used the high-throughput fitness dataset and the stability dataset S2648 (Methods 4.1) to train two different versions of DVE-stability. As shown in Fig.2, using stability changes as the criterion for dividing beneficial mutations (increased stability) and harmful mutations (reduced stability), the The dimensionality reduction results of CLS and ENV features extracted by the two different versions of DVE-stability were compared with the original ESM-2 650M model. On the S669 and Myoglobin test sets, as expected, the original ESM-2 650M model had almost no ability to distinguish between beneficial and harmful mutations. Trained with only fitness data (the high-throughput dataset), DVE-stability demonstrated a certain degree of discrimination for CLS and ENV features labeled with protein stability change, which provided evidence for the correlation between protein “fitness” and “stability”. That is, protein “stability” refers to a protein’s ability to maintain its structure, while “fitness” reflects how well the protein performs its biological function, with stability often being necessary but not sufficient for high fitness [46]. Moreover, DVE-stability trained with stability change data (S2648 dataset) achieved higher discrimination between the two types of mutations.

**Fig. 2.**
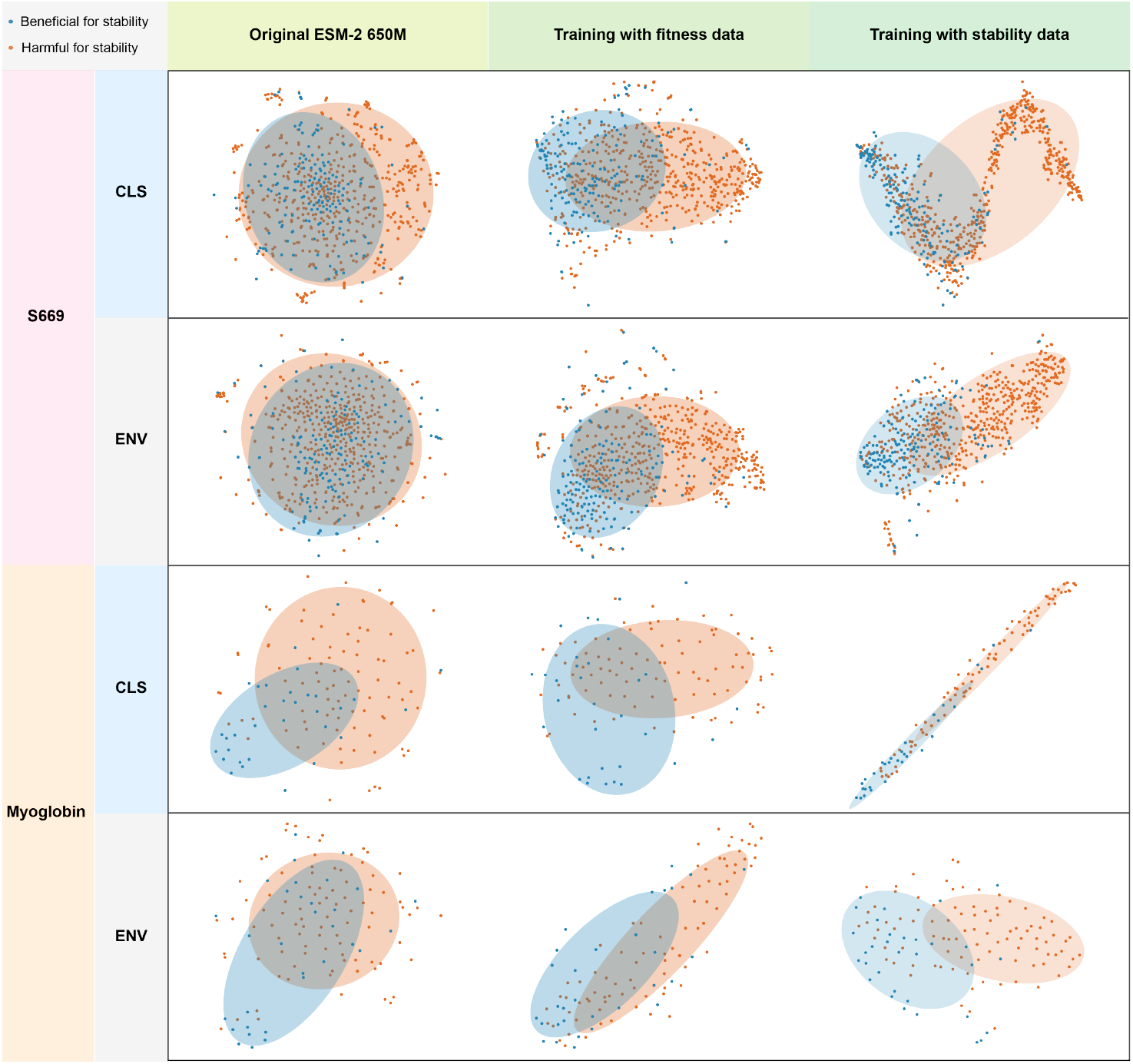
The dimensionality reduction visualization of CLS embedding and ENV embedding of harmful (reduced stability) and beneficial (increased stability) mutations on S669 and Myogloin datasets. **Alt text**: A visualization showing reduced-dimensional embeddings of mutations from S669 and Myoglobin datasets.

### 2.6 Interpretability of DVE-stability

To demonstrate the interpretability of DVE-stability for intramolecular interactions, representative protein G-B1 domain [47] is adopted for case analysis. As shown in Fig.3a, with F52 as the central residue, the top 10 residues with the highest attention scores assigned by DVE-stability model are highlighted. We can find that DVE-stability assigns higher attention scores to some residues (Y3, L5, F30, W43, and Y45) that form *π*™ *π* interactions with F52, which is consistent with the involvement of F52 and these residues in the nucleation event of hydrophobic core [48, 49]. In addition, several enzyme-engineering cases with improved thermal stability are adopted to demonstrate the sensitivity of DVE-stability to fitness effects caused by pairwise mutations. Specifically, the attention scores between pairwise mutations from four representative cases are calculated and visualized, including S238F/W159H on top of *Is*PETase [50], ThermoPETase [51], FAST-PETase [52] and TS-PETase [53]. If the calculated attention score has increased relative to pairwise wild-type residues, we mark it as “positive” with corresponding improvement ratio, otherwise, it is marked as “negative” with corresponding decay ratio. As illustrated in Fig.3b, more than 83% of pairwise mutations exhibit increasing attention scores and only 2 pairs of mutations have decreased attention scores, demonstrating that DVE-stability is sensitive to mutation-induced fitness effects behind the improvement of eznyme thermal stability.

**Fig. 3.**
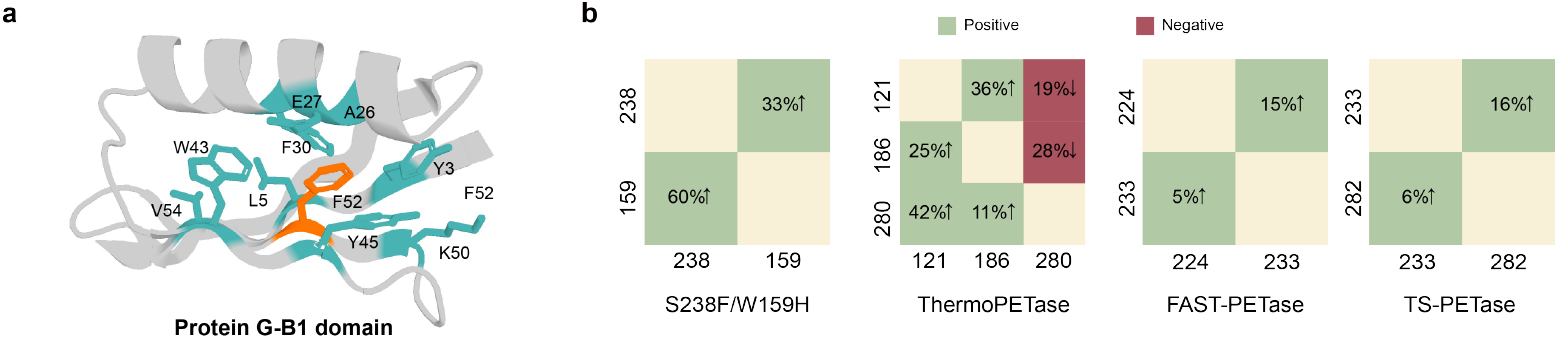
Interpretability of DVE-stability. **a**, The case analysis of representative protein G-B1 domain, where the top 10 residues with the highest attention scores with F52 as the central residue are highlighted. **b**, The attention scores between pairwise mutations from four representative enzyme-engineering cases, where the increased attention scores relative to pairwise wild-type residues are marked as “positive” with corresponding improvement ratio, and the decreased scores are marked as “negative” with corresponding decay ratio. **Alt text**: A two-panel schematic showing attention-based interpretability of DVE-stability on a protein domain and enzyme mutation pairs.

## 3 Discussion

With dual-view ensemble learning, DVE-stability fuses the global interaction dependencies and local interaction dependencies of mutations to predict protein stability changes upon mutations from single sequence. For single-point mutation task scenario, DVE-stability achieved state-of-the-art prediction performance on all 7 benchmark datasets, and comprehensively surpassed other predictive methods on 5 of them. For multiple-point mutation task scenario, DVE-stability outperformed other methods comprehensively through zero-shot behavior. For rare beneficial mutation task scenario, DVE-stability also exhibited superior generalization performance. In addition, DVE-stability is also interpretable in identifying important intramolecular interactions via attention scores. However, DVE-stability still has limitations in predicting unequal-length mutation ((insertion or deletion of amino acids)) tasks. Next, we plan to set special tokens in DVE-stability framework to represent inserted or deleted residues, thereby supporting the prediction of unequal-length mutational stability changes. Looking to the future, as the available data in the community gradually grows, DVE-stability is expected to further improve its prediction performance by training on larger datasets.

## 4 Methods

### 4.1 Dataset

#### 4.1.1 Hypothetical reverse mutation

Protein stability changes can be measured by the difference in folding free energies (ΔΔ*G*) between wild-type protein and their mutants. Considering that the free energy is a thermodynamic state function, ΔΔ*G* is determined by the two endpoints of wild-type protein and mutant, i.e., path independent. Thus, the change of folding free energies from the mutant to the wild-type protein (M → W) ΔΔ*G*_*M→W*_ is the opposite of ΔΔ*G*_*W →M*_, the change of folding free energies from the wild-type to the mutant (W → M):

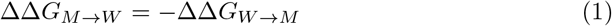

In this work, a reverse dataset composed of hypothetical reverse mutations was created for the benchmark dataset S669 (named S669 reverse). This reverse dataset is adopted to evaluate the predictive symmetry of different methods (i.e., the predicted results of direct and reverse mutations are of opposite signs), which is an aspect of model robustness.

#### 4.1.2 Training datasets

##### High-throughput dataset

The high-throughput dataset was curated from cDNA proteolysis dataset [54] of mega-scale folding stability estimation, which is a protein mutation-induced fitness dataset based on deep mutational scanning. After filtering protein sequences with the sequence identity cutoff of 30% to test dataset using the mmseqs2 library [55] (i.e., training proteins had at most 30% sequence similarity with those in test set), 213,000 mutations (from 97,000 double mutants and 117,000 single mutants) from 116 proteins were adopted by us.

##### S2648 dataset

The S2648 dataset [56] contained 2,648 non-redundant single-point mutants with corresponding ΔΔ*G* obtained from 131 proteins, which has been widely used as a training set for previous stability change prediction methods. The sequences of S2648 had at most 25% sequence similarity with those in its test dataset S669.

##### S8754 dataset

The S8754 dataset [21] contained 8,754 single-point mutations of 301 proteins, which was collected from two ΔΔ*G* resources, ProThermDB [57] and ThermoMutDB [58]. Non-overlapping data were merged and overlapping parts were deduplicated to ensure the uniqueness and high credibility of each data. Sequences with sequence identity greater than 25% with those of the test set (S669 dataset) were removed.

##### C2878 dataset

The C2878 dataset [20] was curated from Q1744 [17], O2567 [11], S2648 [56], and FireProtDB [59]. The duplicates were addressed by taking the ΔΔ*G* measurement with the highest absolute value, and the sequences were filtered by setting a 30% sequence identity cutoff to its corresponding test set (T2837 dataset).

#### 4.1.3 Test datasets

##### S669 dataset

The S669 dataset [9] is a manually curated dataset containing 669 single-point mutants and their ΔΔ*G* measurements (derived from 94 proteins selected from ThermoMutDB [58]), with less than 25% sequence identity to the most used training dataset S2648. This dataset has been used in a large number of previous studies and is a widely-recognized benchmark dataset.

##### S461 dataset

The S461 dataset [60] is a cleaner subset of S669 (461 single-point mutations), obtained by removing erroneous entities in the original S669 dataset.

##### S783 dataset

The S783 dataset [21] was curated from S921 dataset [61], a recognized test dataset for predictive models trained on S2648. The redundant entries were removed, which are either entities that are exactly the same in datasets S921 and S2648 or entities in S921 that are simply the reverse mutations to those in S2648.

##### S349 dataset

The S349 dataset [62] a commonly used blind test set selected from S2648, including 349 single-point mutations.

##### Myoglobin dataset

The Myoglobin dataset [63, 64] is a dataset consisting of 134 single-point mutations in cytoplasmic globular proteins that regulate cellular oxygen concentration.

##### PTEN&TPMT dataset

The PTEN&TPMT dataset [65, 66] is a deep mutational scanning dataset of the phosphatase and tensin homolog (PTEN) and thiopurine S-methyl transferase (TPMT) proteins, with the size of 7,363 single-point mutations.

##### T2837 dataset

The T2837 dataset [20] was curated from P53 [17], Myoglobin, SSym [17], S669, and C5266 [20]. The duplicates were addressed by taking the ΔΔ*G* measurement with the highest absolute value.

##### M1261 dataset

The M1261 dataset [21] is a benchmark dataset for multiple-point mutations. It contained 792 pieces of double mutation data and 469 pieces of triple or higher-order mutation data from 133 proteins.

### 4.2 Architecture overview

Amino acid has 20 different types, represented by a set *AA* = {*A, C*, …, *Y*}. A protein of length *L* can be defined as *S* = (*s*^1^, *s*^2^, …, *s*^*L*^) where *s*^*l*^ ∈ *AA*. Given the position of mutation pos_*mut*_ and a mutated residue 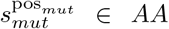 substituting the wild-type residue 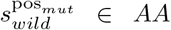, we can get the wild-type protein 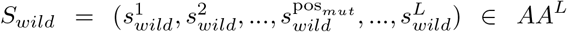 and mutated protein 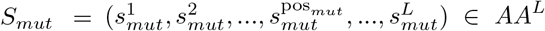. The target of predicting protein stability changes upon point mutation is to calculate the mutation-induced stability change ΔΔ*G*.

DVE-stability is built on the protein language model, ESM-2-650M, that accepts inputs of both wild-type and mutated protein sequences. Both the wild-type and mutated protein sequences, denoted as *S*_*wild*_ and *S*_*mut*_, are input into a shared ESM-2-650M model to generate the corresponding sequence features:

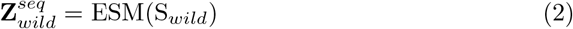

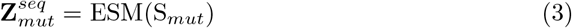

where 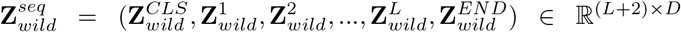 and 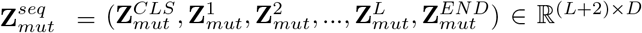 represent the features of wild-type and mutated protein sequences output by ESM-2-650M model, in which *D* represents the dimension of the features.

Subsequently, the sequence features are fed into the structural microenvironment simulation (SMS) module to obtain the local dependencies of mutations as follows:

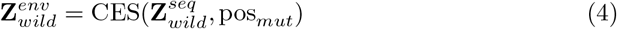

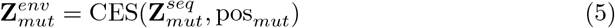

where 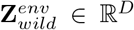 and 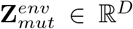 represent the local dependencies of wild-type residue and mutated residue.

Finally, 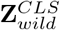 and 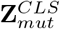 are fed into a ΔΔ*G* prediction neural network, while 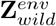 and 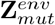 are fed into another parallel ΔΔ*G* prediction neural network. The final predicted ΔΔ*G* is the average of the above two predictions.

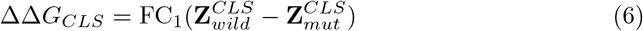

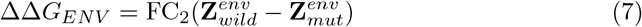

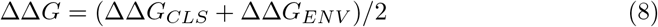

where FC_1_ and FC_2_ adopt the same configuration, that is, a single fully connected layer, to predict the ΔΔ*G*.

### 4.3 SMS module implementation

Given a protein sequence *S*, we obtain its sequence feature **Z**^*seq*^ by feeding *S* into the ESM-2-650M model. Moreover, ESM-2-650M outputs an attention map every transformer layer, where the attention scores reflect the importance of the relationship between the paired tokens. Therefore, we select the structurally adjacent residues based on the attention map *A*^*last*^ ∈ ℝ^*H×*(*L*+2)*×*(*L*+2)^ of the last transformer layer, in which *H* is the number of attention heads of a transformer.

First, *A*^*last*^ is pooled as follows:

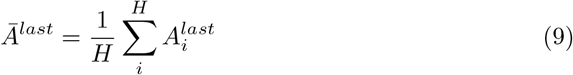

where 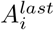 denotes the *i*-th head’s attention map of the *A*^*last*^.

Subsequently, given the mutation position pos_*mut*_, we obtain the attention score vector *A*_*mut*_ ∈ ℝ^*L*+2^ that represents the attention scores to the mutation token (abbreviated as MUT token) by indexing 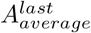 with mutation position pos_*mut*_:

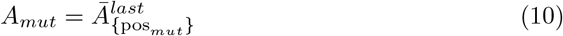

Based on *A*_*mut*_, we select 3 tokens related to MUT token, which rank among the top 3 in attention scores:

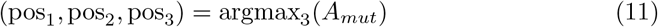

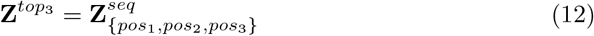

where argmax_3_(·) represents the function to get the 3 most largest elements’ positions of the given vector, and 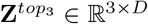 represents the 3 most important token related to MUT token.

Finally, we feed the 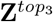 with a learnable embedding vector **Z**^*CLS*^ ∈ ℝ^*D*^ into a one-layer transformer block to integrate them into the final “ENV” embedding of local dependencies.

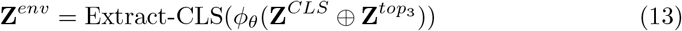

where **Z**^*env*^ ∈ ℝ^*D*^ represents the feature of local dependencies, Extract-CLS denotes extracting the CLS token’s embedding, *ϕ*_*θ*_ represents a one-layer transformer block with trainable parameter *θ*, and ⊕ denotes concatenating the two features.

### 4.4 Attention score calculation of pairwise mutations

In this work, four enzyme-engineering cases were employed to calculate attention scores between pairwise mutations. Specifically, if there are beneficial mutations with mutation positions *P* = {pos_1_, pos_2_, …, pos_*m*_}, we will calculate the difference of attention scores of any two positions (*i, j*) ∈ *P* × *P* between pairwise mutations and pairwise wild-type residues:

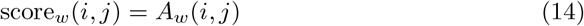

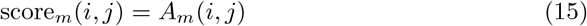

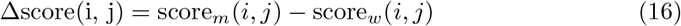

where *A*_*w*_ is the pooled attention map outputted by the last transformer block of ESM-2-650M of pairwise wild-type residues and *A*_*m*_ is that of pairwise mutations.

### 4.5 Evaluation metrics

#### 4.5.1 Mean Absolute Error (MAE)

Mean Absolute Error (MAE) is a metric that measures the average magnitude of errors between predicted and actual values, without considering their direction. It is calculated as the mean of the absolute differences between predictions and true values. A lower MAE indicates better predictive accuracy.

#### 4.5.2 Root mean square error (RMSE)

RMSE is a standard way to measure how closely the predicted measurements align with the true measurements. The formula for RMSE is given by:

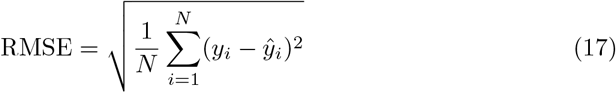

where N is the number of observations, *y*_*i*_ is the actual value of an observation, and ŷ_*i*_ is the predicted value.

#### 4.5.3 Pearson correlation coefficient (*r*)

Pearson correlation coefficient (*r*) is a usual way to measure the linear correlation between two variables, offering a comprehensive analysis when the data distribution permits:

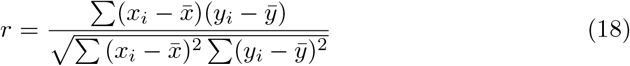

where *x*_*i*_ and *y*_*i*_ are the individual sample points indexed with *i*, 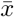 and 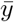 are the sample means.

#### 4.5.4 Spearman correlation coefficient (*ρ*)

The Spearman correlation coefficient (*ρ*) is a non-parametric measure of rank correlation that assesses how well the relationship between two variables can be described by a monotonic function. It evaluates the strength and direction of association between the ranked values of the variables, rather than their raw values.

#### 4.5.5 Kendall correlation coefficient (*τ*)

The Kendall correlation coefficient (*τ*) is a non-parametric measure of rank correlation that evaluates the strength and direction of association between two variables based on the number of concordant and discordant pairs. It reflects the probability that the ranks of two variables are in the same order.

#### 4.5.6 Recall

Recall is the proportion of actual positive cases that are correctly identified by the model. The formula for recall is given by:

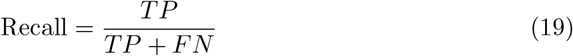

where TP is the number of true positives, TN is the number of true negatives, FP is the number of false positives, and FN is the number of false negatives.

#### 4.5.7 Area Under the Receiver Operating Characteristic Curve (AUC)

The Area Under the Receiver Operating Characteristic Curve (AUC) measures a model’s ability to distinguish between classes across all possible classification thresholds. It represents the probability that a randomly chosen positive instance is ranked higher than a randomly chosen negative one.

#### 4.5.8 Matthews Correlation Coefficient (MCC)

The Matthews Correlation Coefficient (MCC) is a balanced metric that evaluates the quality of binary classifications, taking into account true and false positives and negatives. It is especially useful for imbalanced datasets, as it considers all four outcomes in the confusion matrix.

#### 4.5.9 Top-K precision

Top-K precision [29] measures the proportion of true top K mutations correctly identified among the predicted top K, serving as an indicator of success rate in practical protein engineering applications.

## Key Points

- We propose a dual-view ensemble learning-based framework, DVE-stability, for mutation-induced protein stability change prediction from single sequence.
- DVE-stability integrates the global dependencies of mutations dominated by longrange interactions and local dependencies of mutations determined by the structural microenvironments, enabling the neural network to capture the intramolecular interaction network in which the mutation is located from two views through ensemble learning.
- A structural microenvironment simulation (SMS) module is designed to indirectly introduce the information of structural microenvironment at the sequence level.
- DVE-stability achieved state-of-the-art prediction performance on 7 single-point mutation benchmark datasets, and demonstrated superior generalization performance for multiple-point mutations and rare beneficial mutations.
- DVE-stability represents an interpretable ensemble learning approach for protein stability change prediction using only sequence, which is expected to provide a flexible and efficient tool for practical protein engineering.

## Biographical note

Dr Chen’s laboratory is committed to promoting the intersection of state-of-the-art deep-learning technologies and life sciences, focusing on foundation models in the field of life sciences and their applications to proteins and small molecules.

## Author contributions

Zhiwei Nie and Yiming Ma conceived the concept; Zhiwei Nie and Yiming Ma designed the deep-learning framework, trained models, and performed experiments. Yutian Liu, Xiansong Huang, Zhihong Liu, Peng Yang, Fan Xu, Feng Yin, and Zigang Li collected the data involved in the experiments and provided helpful discussions. Jie Chen, Wen-Bin Zhang, Zhixiang Ren, and Jie Fu supervised this research and provided helpful discussions. All authors contributed to the writing and revision of the manuscript.

## Funding

This work was supported in part by the Shenzhen Medical Research Funds in China (No. B2302037), Natural Science Foundation of China (No. 61972217, 32071459, 62176249, 62006133, 62271465), Guangdong S&T Program (Grant No. 2024B0101010003), and AI for Science (AI4S)-Preferred Program, Peking University Shenzhen Graduate School, China.

## Competing interests

The authors declare no competing interests.

## Data and code availability

Relevant data, code and models are available via GitHub at https://github.com/ZhiweiNiepku/DVE-stability.

## Notes

### Competing Interest Statement

The authors have declared no competing interest.

### Summary of Updates

We have updated the manuscript, including figures, text, etc.

## References

[1] Reva, B., Antipin, Y. & Sander, C. Predicting the functional impact of protein mutations: application to cancer genomics. Nucleic acids research 39, e118–e118 (2011).

[2] Wylie, C. S. & Shakhnovich, E. I. A biophysical protein folding model accounts for most mutational fitness effects in viruses. Proceedings of the National Academy of Sciences 108, 9916–9921 (2011).

[3] Tokuriki, N. & Tawfik, D. S. Stability effects of mutations and protein evolvability. Current opinion in structural biology 19, 596–604 (2009).

[4] Pucci, F., Bernaerts, K. V., Kwasigroch, J. M. & Rooman, M. Quantification of biases in predictions of protein stability changes upon mutations. Bioinformatics 34, 3659–3665 (2018).

[5] Wu, S., Snajdrova, R., Moore, J. C., Baldenius, K. & Bornscheuer, U. T. Biocatalysis: enzymatic synthesis for industrial applications. Angewandte Chemie International Edition 60, 88–119 (2021).

[6] Bell, E. L. et al. Biocatalysis. Nature Reviews Methods Primers 1, 1–21 (2021).

[7] Jay, S. M. & Lee, R. T. Protein engineering for cardiovascular therapeutics: untapped potential for cardiac repair. Circulation research 113, 933–943 (2013).

[8] Meghwanshi, G. K. et al. Enzymes for pharmaceutical and therapeutic applications. Biotechnology and applied biochemistry 67, 586–601 (2020).

[9] Pancotti, C. et al. Predicting protein stability changes upon single-point mutation: a thorough comparison of the available tools on a new dataset. Briefings in Bioinformatics 23, bbab555 (2022).

[10] Pucci, F., Schwersensky, M. & Rooman, M. Artificial intelligence challenges for predicting the impact of mutations on protein stability. Current opinion in structural biology 72, 161–168 (2022).

[11] Caldararu, O., Mehra, R., Blundell, T. L. & Kepp, K. P. Systematic investigation of the data set dependency of protein stability predictors. Journal of Chemical Information and Modeling 60, 4772–4784 (2020).

[12] Qiu, Y., Huang, T. & Cai, Y.-D. Review of predicting protein stability changes upon variations. Proteomics 2300371 (2024).

[13] Schymkowitz, J. et al. The foldx web server: an online force field. Nucleic acids research 33, W382–W388 (2005).

[14] Sheffler, W. & Baker, D. Rosettaholes: rapid assessment of protein core packing for structure prediction, refinement, design, and validation. Protein Science 18, 229–239 (2009).

[15] Zhou, H. & Zhou, Y. Distance-scaled, finite ideal-gas reference state improves structure-derived potentials of mean force for structure selection and stability prediction. Protein science 11, 2714–2726 (2002).

[16] Pires, D. E., Ascher, D. B. & Blundell, T. L. mcsm: predicting the effects of mutations in proteins using graph-based signatures. Bioinformatics 30, 335–342 (2014).

[17] Li, B., Yang, Y. T., Capra, J. A. & Gerstein, M. B. Predicting changes in protein thermodynamic stability upon point mutation with deep 3d convolutional neural networks. PLoS computational biology 16, e1008291 (2020).

[18] Savojardo, C., Fariselli, P., Martelli, P. L. & Casadio, R. Inps-md: a web server to predict stability of protein variants from sequence and structure. Bioinformatics 32, 2542–2544 (2016).

[19] Dieckhaus, H., Brocidiacono, M., Randolph, N. Z. & Kuhlman, B. Transfer learning to leverage larger datasets for improved prediction of protein stability changes. Proceedings of the national academy of sciences 121, e2314853121 (2024).

[20] Diaz, D. J. et al. Stability oracle: a structure-based graph-transformer framework for identifying stabilizing mutations. Nature Communications 15, 6170 (2024).

[21] Xu, Y., Liu, D. & Gong, H. Improving the prediction of protein stability changes upon mutations by geometric learning and a pre-training strategy. Nature Computational Science 1–11 (2024).

[22] Dauparas, J. et al. Robust deep learning–based protein sequence design using proteinmpnn. Science 378, 49–56 (2022).

[23] Pancotti, C. et al. A deep-learning sequence-based method to predict protein stability changes upon genetic variations. Genes 12, 911 (2021).

[24] Folkman, L., Stantic, B., Sattar, A. & Zhou, Y. Ease-mm: sequence-based prediction of mutation-induced stability changes with feature-based multiple models. Journal of molecular biology 428, 1394–1405 (2016).

[25] Montanucci, L., Capriotti, E., Frank, Y., Ben-Tal, N. & Fariselli, P. Ddgun: an untrained method for the prediction of protein stability changes upon single and multiple point variations. BMC bioinformatics 20, 1–10 (2019).

[26] Cheng, J., Randall, A. & Baldi, P. Prediction of protein stability changes for single-site mutations using support vector machines. Proteins: Structure, Function, and Bioinformatics 62, 1125–1132 (2006).

[27] Umerenkov, D. et al. Prostata: a framework for protein stability assessment using transformers. Bioinformatics 39, btad671 (2023).

[28] Ouyang-Zhang, J., Diaz, D., Klivans, A. & Krähenbühl, P. Predicting a protein’s stability under a million mutations. Advances in Neural Information Processing Systems 36 (2024).

[29] Chen, Y., Xu, Y., Liu, D., Xing, Y. & Gong, H. An end-to-end framework for the prediction of protein structure and fitness from single sequence. Nature Communications 15, 7400 (2024).

[30] Lin, Z. et al. Evolutionary-scale prediction of atomic-level protein structure with a language model. Science 379, 1123–1130 (2023).

[31] Jumper, J. et al. Highly accurate protein structure prediction with alphafold. Nature 596, 583–589 (2021).

[32] Gillioz, A., Casas, J., Mugellini, E. & Abou Khaled, O. Overview of the transformer-based models for nlp tasks, 179–183 (IEEE, 2020).

[33] Sun, C., Qiu, X., Xu, Y. & Huang, X. How to fine-tune bert for text classification?, 194–206 (Springer, 2019).

[34] Liu, T., Wang, X., Lv, C., Zhen, R. & Fu, G. Sentence matching with syntax-and semantics-aware bert, 3302–3312 (2020).

[35] Kim, T., Yoo, K. M. & Lee, S.-g. Self-guided contrastive learning for bert sentence representations, 2528–2540 (2021).

[36] Shroff, R. et al. Discovery of novel gain-of-function mutations guided by structurebased deep learning. ACS synthetic biology 9, 2927–2935 (2020).

[37] Lu, H. et al. Machine learning-aided engineering of hydrolases for pet depolymerization. Nature 604, 662–667 (2022).

[38] d’Oelsnitz, S. et al. Biosensor and machine learning-aided engineering of an amaryllidaceae enzyme. Nature Communications 15, 2084 (2024).

[39] Vig, J. et al. Bertology meets biology: Interpreting attention in protein language models (2021).

[40] Rives, A. et al. Biological structure and function emerge from scaling unsupervised learning to 250 million protein sequences. Proceedings of the National Academy of Sciences 118, e2016239118 (2021).

[41] Rodrigues, C. H., Pires, D. E. & Ascher, D. B. Dynamut2: Assessing changes in stability and flexibility upon single and multiple point missense mutations. Protein Science 30, 60–69 (2021).

[42] Zhou, Y., Pan, Q., Pires, D. E., Rodrigues, C. H. & Ascher, D. B. Ddmut: predicting effects of mutations on protein stability using deep learning. Nucleic Acids Research 51, W122–W128 (2023).

[43] Arnold, F. H. Directed evolution: bringing new chemistry to life. Angewandte Chemie (International Ed. in English) 57, 4143 (2017).

[44] Hie, B. L. et al. Efficient evolution of human antibodies from general protein language models. Nature Biotechnology 42, 275–283 (2024).

[45] Cui, Y. et al. Computational redesign of a petase for plastic biodegradation under ambient condition by the grape strategy. Acs Catalysis 11, 1340–1350 (2021).

[46] Tokuriki, N., Stricher, F., Serrano, L. & Tawfik, D. S. How protein stability and new functions trade off. PLoS computational biology 4, e1000002 (2008).

[47] Sjöbring, U., Björck, L. & Kastern, W. Streptococcal protein g. gene structure and protein binding properties. Journal of biological chemistry 266, 399–405 (1991).

[48] Kmiecik, S. & Kolinski, A. Folding pathway of the b1 domain of protein g explored by multiscale modeling. Biophysical journal 94, 726–736 (2008).

[49] Blanco, F. J. & Serrano, L. Folding of protein g b1 domain studied by the conformational characterization of fragments comprising its secondary structure elements. European journal of biochemistry 230, 634–649 (1995).

[50] Austin, H. P. et al. Characterization and engineering of a plastic-degrading aromatic polyesterase. Proceedings of the National Academy of Sciences 115, E4350–E4357 (2018).

[51] Son, H. F. et al. Rational protein engineering of thermo-stable petase from ideonella sakaiensis for highly efficient pet degradation. Acs Catalysis 9, 3519–3526 (2019).

[52] Lu, H. et al. Deep learning redesign of petase for practical pet degrading applications. Biorxiv 2021–10 (2021).

[53] Zhong-Johnson, E. Z. L., Voigt, C. A. & Sinskey, A. J. An absorbance method for analysis of enzymatic degradation kinetics of poly (ethylene terephthalate) films. Scientific Reports 11, 928 (2021).

[54] Tsuboyama, K. et al. Mega-scale experimental analysis of protein folding stability in biology and design. Nature 620, 434–444 (2023).

[55] Steinegger, M. & Söding, J. Mmseqs2 enables sensitive protein sequence searching for the analysis of massive data sets. Nature biotechnology 35, 1026–1028 (2017).

[56] Dehouck, Y. et al. Fast and accurate predictions of protein stability changes upon mutations using statistical potentials and neural networks: Popmusic-2.0. Bioinformatics 25, 2537–2543 (2009).

[57] Nikam, R., Kulandaisamy, A., Harini, K., Sharma, D. & Gromiha, M. M. Prothermdb: thermodynamic database for proteins and mutants revisited after 15 years. Nucleic acids research 49, D420–D424 (2021).

[58] Xavier, J. S. et al. Thermomutdb: a thermodynamic database for missense mutations. Nucleic acids research 49, D475–D479 (2021).

[59] Stourac, J. et al. Fireprotdb: database of manually curated protein stability data. Nucleic acids research 49, D319–D324 (2021).

[60] Hernández, I. M., Dehouck, Y., Bastolla, U., López-Blanco, J. R. & Chacón, P. Predicting protein stability changes upon mutation using a simple orientational potential. Bioinformatics 39, btad011 (2023).

[61] Chen, Y. et al. Premps: Predicting the impact of missense mutations on protein stability. PLoS computational biology 16, e1008543 (2020).

[62] Iqbal, S. et al. Prost: Alphafold2-aware sequence-based predictor to estimate protein stability changes upon missense mutations. Journal of chemical information and modeling 62, 4270–4282 (2022).

[63] Ordway, G. A. & Garry, D. J. Myoglobin: an essential hemoprotein in striated muscle. Journal of Experimental Biology 207, 3441–3446 (2004).

[64] Wang, S., Tang, H., Shan, P., Wu, Z. & Zuo, L. Pros-gnn: predicting effects of mutations on protein stability using graph neural networks. Computational Biology and Chemistry 107, 107952 (2023).

[65] Lv, X. et al. Accurately predicting mutation-caused stability changes from protein sequences using extreme gradient boosting. Journal of chemical information and modeling 60, 2388–2395 (2020).

[66] Li, G., Yao, S. & Fan, L. Prostage: Predicting effects of mutations on protein stability by using protein embeddings and graph convolutional networks. Journal of Chemical Information and Modeling 64, 340–347 (2024).

